# Multiscale, Survey-and-Target Optical Super-Resolution Histology

**DOI:** 10.64898/2026.10.26.762070

**Authors:** Kidan Tadesse, Oliver Lee, Hajun Yoo, Ronghao Zhang, Qiaoni Liu, Peng Chen, Zizhen Zha, Gabriel A. Kwong, Vaunita Parihar, Jennifer Robertson, Brian Robinson, Eunice Chou, Shu Jia

## Abstract

Super-resolution microscopy reveals fine subcellular tissue architecture, yet extending this capability to histological screening requires balancing spatial resolution with imaging area, axial access, throughput, and instrumental complexity. Here we present multiscale optical super-resolution histology (MOSH), an integrated optical–computational platform that combines tissue-scale, extended-depth-of-focus super-resolution surveys with coordinate-guided three-dimensional interrogation of selected targets. MOSH leverages stationary multifocal excitation, continuous sample translation, tunable axial refocusing, and parallelized reconstruction within a shared spatial-coordinate framework in a conventional epi-fluorescence setting. This survey-and-target strategy preserves tissue-scale context, directs volumetric acquisition to selected subcellular structures, and generates virtual H&E images that retain resolution-enhanced structural information. Across cultured cells, cytological preparations, conventional tissue sections, and intact unsectioned biopsies, MOSH achieved approximately twofold resolution enhancement, continuous lateral coverage beyond the native field of view, and tunable axial survey ranges up to 20 µm. Bridging tissue-scale histological screening and targeted, quantitative nanoscale interrogation, MOSH establishes a scalable framework for multiscale analysis of biological and clinical specimens.

## INTRODUCTION

Optical microscopy underpins histological assessment, traditionally through the examination of physically sectioned and stained specimens. Fluorescence microscopy extends this capability by enabling rapid, sensitive visualization of molecularly defined structures within heterogeneous tissue microenvironments(*1–4*). When coupled with optical sectioning, fluorescence imaging can recover volumetric information from intact or minimally processed specimens, allowing nuclear and extranuclear features to be examined within their three-dimensional context(*5–7*). These capabilities are increasingly being translated into tissue assessment, including, for example, autofluorescence-based virtual staining that transforms label-free signals into histology-like images(*8, 9*), slide-free fluorescence methods that enable rapid, non-destructive evaluation of intact specimens(*10, 11*), and fluorescent analogues of hematoxylin and eosin (H&E) that generate high-contrast, pathologist-readable images without conventional chromogenic staining(*12*). For large-area tissue assessment, wide-field architectures are particularly well suited because they combine high imaging throughput with operational simplicity.

Conventional wide-field systems, however, remain constrained by the diffraction limit and therefore cannot resolve nanoscale subcellular features that may contain diagnostically relevant information(13, 14). Super-resolution microscopy (SRM) addresses this barrier by revealing fine structural detail in cells and tissues(15–18). Among these methods, structured illumination microscopy (SIM) is particularly attractive for histological imaging because it combines wide-field camera-based parallel detection, which supports relatively rapid acquisition, efficient photon use, and lower phototoxicity than coordinate-targeted nanoscopy(19–22). Meanwhile, image-scanning microscopy (ISM) provides a complementary, confocal-derived strategy: focused excitation is scanned across the specimen, spatially resolved detection records the emitted light, and pixel reassignment narrows the effective point-spread function while preserving confocal optical sectioning and background rejection(23–26). Although these properties make ISM attractive for optically heterogeneous specimens, SIM-based systems have already been applied to rapid, large-area imaging of tissue sections and clinical specimens(27–30).

Despite this progress, current SRM platforms rarely combine the spatial resolution, imaging area, axial access, and operational simplicity required for routine histological workflows(14, 31). Large-area screening generally relies on low-numerical-aperture objectives to obtain broad fields of view and extended depth of focus, but at the expense of spatial resolution. Conversely, high-numerical-aperture systems typically cover comparatively small fields of view over shallow native depths(32, 33). Conventional SIM requires multiple phase- and orientation-shifted acquisitions, and illumination-pattern contrast can deteriorate in thick or strongly scattering tissue(20, 34). Point-scanning ISM effectively rejects out-of-focus background(35), but sequential raster scanning restricts its throughput. Both modalities may additionally require specialized hardware, precise calibration and alignment, and computationally intensive reconstruction(36–38). The unmet need is therefore not resolution alone, but an imaging architecture that integrates tissue-scale survey imaging with targeted volumetric analysis at subdiffractional resolution while maintaining high throughput and straightforward operation.

Here, we introduce multiscale optical super-resolution histology (MOSH), an integrated optical– computational framework for tissue-scale survey imaging and targeted subcellular analysis. In a conventional epi-fluorescence setting, MOSH leverages stationary multifocal excitation, continuous sample translation, rapid axial refocusing, and spatial registration to support two complementary modes: high-throughput, super-resolved extended-depth-of-focus survey imaging and targeted three-dimensional super-resolution imaging of selected regions of interest. A shared coordinate framework connects the reconstructed survey image to the physical specimen, enabling selected structures to be relocated and examined volumetrically without transferring the sample or changing imaging systems. A parallelized computational pipeline reconstructs continuous data streams, segments and ranks nuclei, supports user-directed target selection, and converts dual-channel fluorescence data into virtual H&E images. We validate MOSH using cultured cells, cytological preparations, conventional tissue sections, and fresh unsectioned biopsies, demonstrating twofold resolution enhancement across extended lateral areas and axial ranges. By integrating acquisition, reconstruction, target selection, and coordinate-aware reimaging into a unified workflow, MOSH establishes a practical survey-and-target framework for multiscale optical histology.

## RESULTS

### The Principle and Framework of MOSH

MOSH integrates tissue-scale survey imaging, subcellular three-dimensional targeted imaging, automated data handling, and virtual H&E rendering on a conventional epifluorescence microscope (**Fig. 1a, Methods, Supplementary Note 1**). The unified optical and spatial-coordinate framework allows the same specimen to be surveyed, computationally analyzed, physically revisited, and interrogated volumetrically within a single workflow (**Supplementary Fig. S1**).

**Fig. 1.**
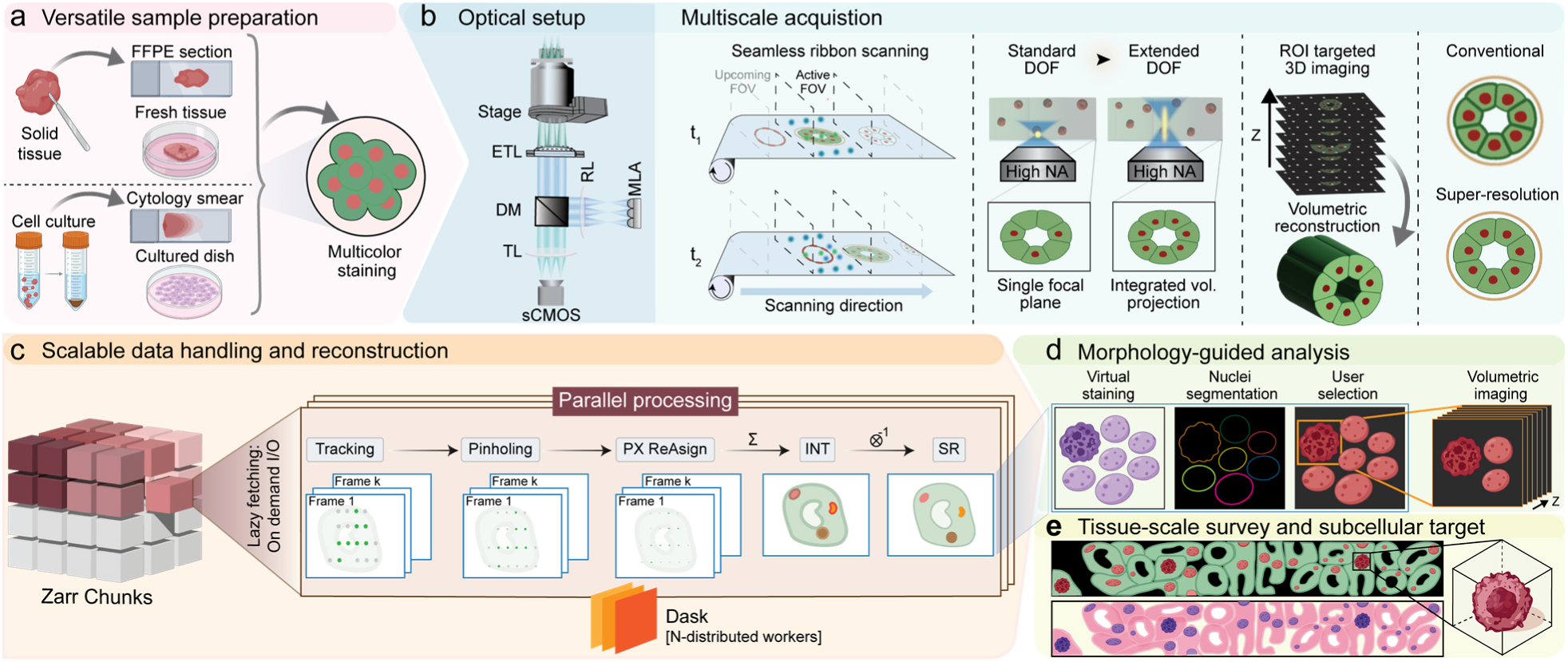
Overview of the multiscale optical super-resolution histology (MOSH) platform. (a) Sample compatibility of MOSH, including sectioned tissue, unsectioned biopsies, and cytological preparations labeled with fluorescent hematoxylin and eosin (H&E) analogues. (b) Optical configuration for multiscale 2D and 3D imaging. Left-to-right insets show continuous multifocal ribbon scanning, electrically tunable lens (ETL)-based extended-depth-of-field (EDoF) imaging, piezoelectric Z-stage-based volumetric acquisition, and 2× resolution enhancements relative to conventional wide-field imaging. (c) Parallelized data-processing and reconstruction pipeline, including Zarr-based data storage, lazy loading, tracking, pinholing, pixel reassignment, signal summation, and deconvolution within a Dask computing framework. (d) AI-assisted nuclear segmentation and morphological atypia scoring with interactive region-of-interest (ROI) selection for targeted 3D imaging. (e) Stitching of reconstructed two-channel super-resolution fluorescence, computational translation into virtual H&E images, and target volumetric imaging.

The optical architecture integrates stationary multifocal excitation generated by a microlens array (MLA), continuous sample translation, high-speed sCMOS detection, and programmable axial control to implement two complementary super-resolution modes: large-area survey imaging and targeted volumetric acquisition (**Fig. 1b, Supplementary Fig. S2, Methods**). In survey mode, MOSH achieves high-content acquisition by integrating tunable electro-optics for rapid axial refocusing with synchronous sample translation, thereby expanding both the accessible depth range and the lateral imaging area beyond the native epi-fluorescence content(*39*) (**Fig. 1b, Methods**). In targeted mode, the axial-control mechanism acquires a dense three-dimensional volume at a selected location within the specimen(*40*) (**Methods**).

In practice, a synchronized acquisition and data-handling pipeline supports continuous image capture without interruption by data transfer or storage (**Supplementary Fig. S1, Supplementary Note 1**). The resulting resolution-doubled survey image provides tissue-scale context and a spatial reference for targeted volumetric interrogation (**Fig. 1c, Methods**). Morphology-guided analysis segments and prioritizes candidate structures for user review, after which a calibrated coordinate transformation maps each selected ROI from image space to its physical position on the specimen (**Fig. 1d, Methods**). The microscope then returns to the selected location and automatically acquires a three-dimensional super-resolution volume, directly connecting the tissue-scale survey image to localized volumetric information (**Fig. 1d, e, Methods**). MOSH further converts the reconstructed nuclear and cytoplasmic fluorescence channels into virtual H&E images, retaining super-resolved structural information within a pathologist-familiar histological representation (**Fig. 1e, Methods**). Together, these capabilities establish MOSH as a unified survey-and-target platform that links large-area super-resolution screening with coordinate-guided volumetric interrogation and virtual histology (**Supplementary Table 1**).

### High-throughput super-resolution screening across diverse cytological preparations

Clinical cytology and hematology require rapid imaging across large specimen areas, yet cellular overlap and uneven mounting topography can compromise high-resolution visualization(*41, 42*). Here, we evaluated MOSH using K562 human erythroleukemia cells prepared in three formats representing distinct cytological conditions (**Fig. 2a, Methods**): cells cultured in a dish to preserve three-dimensional morphology, cells deposited on a coverslip and mounted on a slide to model a wet mount, and centrifuged cells smeared directly onto a slide to model a conventional hematological smear.

**Fig. 2.**
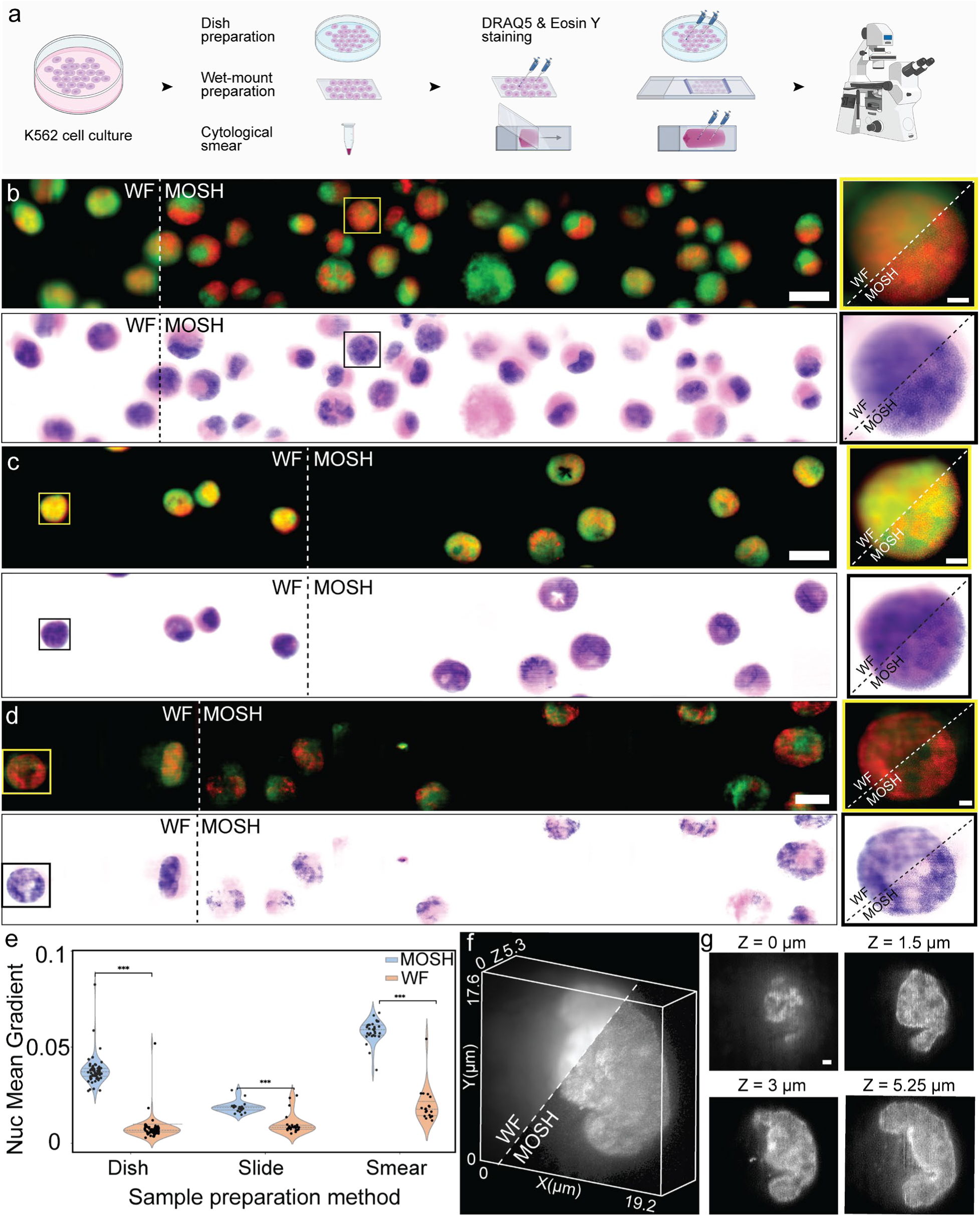
Large-area super-resolution survey and targeted volumetric imaging of K562 cells across cytological preparations. (a) Schematic of K562 sample preparation, fluorescence labeling and MOSH imaging. Cells were prepared by dish culture, coverslip deposition followed by wet mounting, or cytological smearing of a centrifuged cell pellet. (b–d) Stacked wide-field (WF) and MOSH extended-depth-of-focus (EDoF) fluorescence images (top) and corresponding virtual H&E renderings (bottom) of cells prepared by dish culture (b), wet mounting (c), and cytological smearing (d). Green, Eosin Y, labeling cytoplasmic and extracellular structures; DRAQ5, labeling nuclei. Insets: diagonally divided zoomed-in boxed cells in the WF and MOSH images. (e) Mean nuclear intensity gradients measured from the WF and MOSH images for the three preparation formats. Each point represents one nucleus (n = 19–65 nuclei per condition); statistical significance was assessed using two-sided Welch’s t-tests. *P < 0.05, **P < 0.01 and ***P < 0.001. (f) Diagonally divided three-dimensional rendering comparing the WF and MOSH reconstructions of a survey-selected, dish-cultured K562 nucleus. (g) Selected optical sections from the MOSH volume in (f) at the indicated axial positions. MOSH acquired a continuous 1.18 mm × 65 µm survey over a 5-µm EDoF, and representative 420-µm-wide regions are shown in (b–d). Scale bars: 20 µm (b–d) and 3 µm (b–d, insets, and g).

In practice, we demonstrated MOSH for continuous acquisition across a millimeter-scale field of view, with an extended depth of focus of at least 5 µm, adjusted to the thickness of each cytological preparation (**Methods**). Relative to conventional wide-field microscopy, this represents a 1-2 orders-of-magnitude expansion in imaging area and a feasible 10-fold increase in axial coverage of entire cells. Across the three K562 preparation formats, MOSH yielded a lateral resolution of approximately 140 nm based on decorrelation analysis(*43*), an extended focal depth of up to 20 µm, and a survey-dimensional throughput up to approximately 15,320 µm^3^ s⁻¹ (**Supplementary Note 2**), while consistently improving the delineation of nuclear and cytoplasmic boundaries (**Fig. 2b-d, top**). This capability enabled the capture of extensive intranuclear DNA-poor regions, consistent with nucleoli, which were otherwise obscured by low resolution and out-of-focus fluorescence in wide-field scanning projections (**Fig. 2b-d, insets**).

Importantly, MOSH converted the native super-resolution fluorescence data into virtual H&E images that retained these subdiffractional structural features and quantitative contrast, making nanoscale nuclear and cytoplasmic features accessible within a familiar histological representation across all sample-preparation formats (**Fig. 2b-d, bottom**). We quantified the improvement in nuclear structural definition using the mean nuclear intensity gradient (**Supplementary Note 2**). MOSH yielded significantly higher nuclear gradients than wide-field imaging across all three preparation formats (**Fig. 2e**). The largest improvements occurred in the dish and smear conditions, indicating a substantial loss of structural sharpness in the corresponding wide-field images.

MOSH subsequently used the survey image as a spatial reference to return to the mapped location of a dish-cultured K562 cell and reconstruct its intact nucleus throughout its axial extent (**Fig. 2f, g**). Across individual optical sections, the reconstructed volume resolved depth-dependent changes in nuclear morphology, with lateral and axial resolutions of 150 nm and 300 nm(*40*), respectively, over a 5.25-µm axial acquisition range without mechanically flattening the cell (**Supplementary Note 2**). This survey-guided reacquisition directly connected population-scale screening with spatially registered volumetric analysis of an individual target. MOSH further demonstrated large-area, depth-extended super-resolution imaging in a densely packed human blood smear, supporting its applicability to high-content hematological specimens (**Supplementary Fig. S3, Supplementary Note 2**). Collectively, these results establish MOSH as a multiscale cytology platform that preserves population-level context while providing quantitative subcellular contrast and targeted three-dimensional super-resolution analysis of individual cells across diverse preparation formats.

### Survey-and-target super-resolution analysis of human colon cancer tissue

Histopathological assessment of cancer relies on evaluating nuclear morphology, chromatin organization, and glandular architecture across spatially heterogeneous tissue regions (*44–46*). We therefore applied MOSH to patient-derived stage I human colon adenocarcinoma sections (**Fig. 3a, Methods**). In survey mode, MOSH imaged a continuous millimeter-scale area across the full 5-µm section thickness, capturing both pathologist-annotated benign (**Fig. 3b-g**) and malignant (**Fig. 3h-k**) regions. Compared with wide-field imaging, the super-resolution reconstructions improved optical sectioning and delineation of glandular and nuclear structures while preserving the surrounding tissue context (**Fig. 3b, h, Supplementary Fig. S4**). Conversion of the native super-resolution fluorescence data into virtual H&E images transferred this structural information into a familiar histological representation with enhanced optical clarity (**Fig. 3c, i, Supplementary Note 3, Supplementary Fig. S5**). Specifically, MOSH clearly resolved glandular boundaries and internal nuclear features within both benign and malignant tissue. In benign regions, both fluorescence and virtual H&E images revealed sharper epithelial and nuclear boundaries than their wide-field counterparts (**Fig. 3d-g**), exhibiting reduced out-of-focus background and increased modulation depth in dense tissue and individual nuclei. Moreover, MOSH showed irregular glandular organization and heterogeneous intranuclear structures within densely packed tubules in malignant regions, key histopathologic descriptors of malignant behavior that were otherwise substantially obscured by wide-field blur(*46, 47*) (**Fig. 3j, k**).

**Fig. 3.**
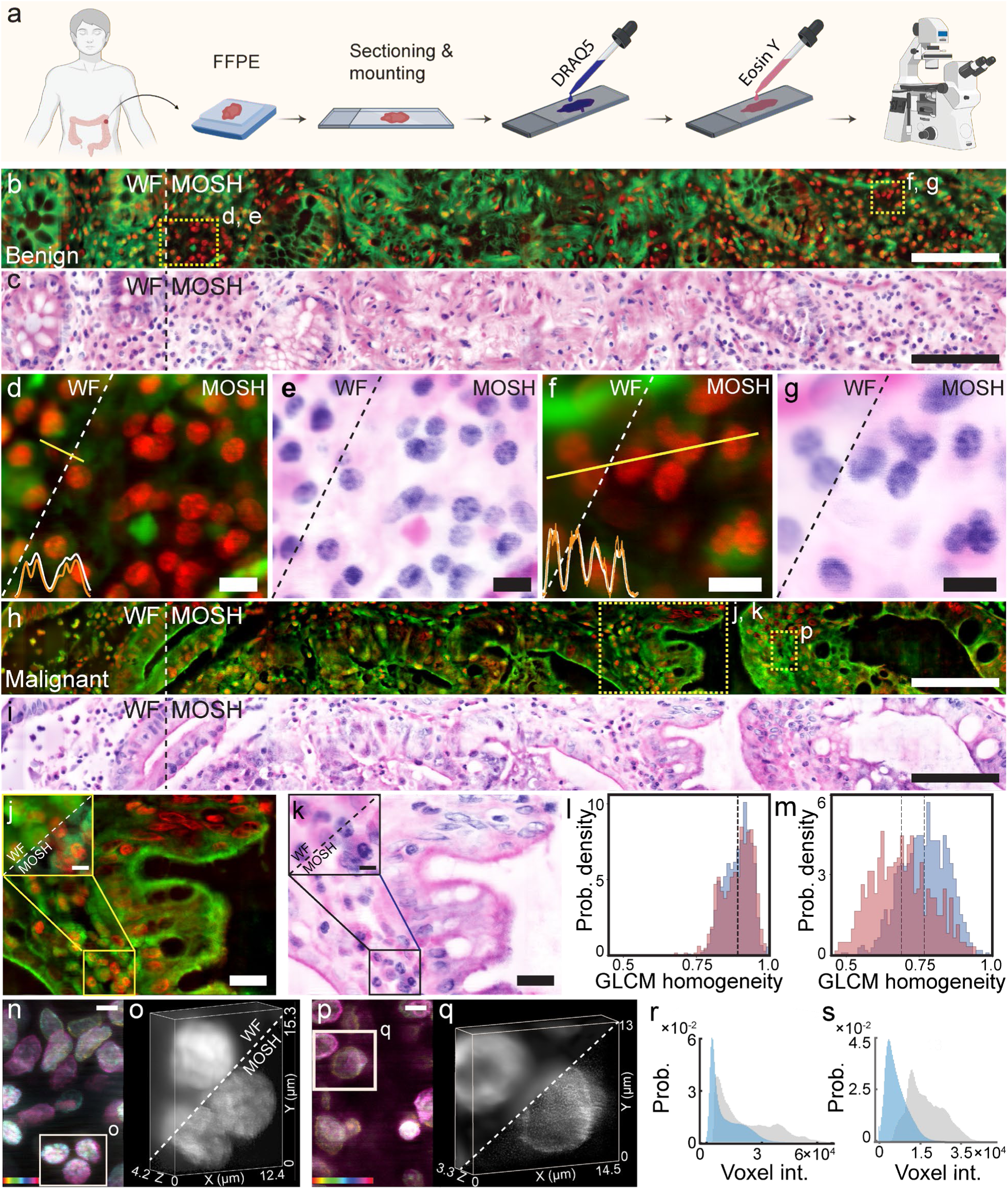
Multiscale survey-and-target MOSH analysis of stage I human colon adenocarcinoma tissue. (a) Schematic of FFPE tissue preparation, sectioning, fluorescence labeling, and MOSH imaging. Red, DRAQ5; green, Eosin Y. (b, c) 5-µm ex-tended-depth-of-focus (EDoF) survey of a pathologist-annotated benign region, shown as fluorescence (b) and correspond-ing virtual H&E (c) images. The vertical dashed line separates the stacked wide-field (WF) and MOSH reconstructions. (d–g) Diagonally divided zoomed-in WF and MOSH images of the boxed regions in (b, c). Line-intensity profiles along the indicated transects compare the spatial modulation recovered by WF and MOSH imaging. (h, i) 5-µm EDoF survey of a pathologist-annotated malignant region, shown as fluorescence (h) and corresponding virtual H&E (i) images. (j, k) Diag-onally divided zoomed-in WF and MOSH fluorescence (j) and virtual H&E (k) images of the boxed region in (h, i). Insets: zoomed-in subregions of the WF and MOSH images. (l, m) Per-nucleus distributions of gray-level co-occurrence matrix (GLCM) homogeneity calculated from intranuclear pixels in the benign (blue) and malignant (red) regions under WF (l) and MOSH (m) imaging. Each value represents one segmented nucleus (WF: n = 927 benign and 569 malignant; MOSH SR: n = 942 benign and 543 malignant); distribution separation was quantified using Cliff’s δ and the area under the receiver operating-characteristic curve. (n, p) Depth-color-coded projections of survey-selected benign (n) and malignant (p) regions, with color (n, 0-4.2 µm; p, 0-3.3 µm) denoting axial position. (o, q) Diagonally divided zoomed-in WF and MOSH volumetric reconstructions of the boxed regions in n and p, respectively. (r, s) Voxel-intensity distributions for the WF (gray) and MOSH (blue) reconstructions of the benign (r) and malignant (s) volumes. MOSH acquired a continuous 1.146 mm × 114 µm field of view in (b, c, h, i). Scale bars: 100 µm (b, c, h, i), 10 µm (d–g), 20 µm (j, k), and 5 µm (j, k, in-sets, and n, p).

We next examined the MOSH-facilitated quantitative analysis of nuclear texture. In practice, nuclei were segmented using identical parameters in corresponding wide-field and super-resolution images, and then gray-level co-occurrence matrix (GLCM) features were derived from their interior pixels(*48–50*) (**Methods, Supplementary Note 4**). Under wide-field imaging, GLCM homogeneity provided essentially no separation between benign and malignant nuclei (Cliff’s δ = −0.02, where 0 indicates complete overlap and positive values denote lower homogeneity in malignant nuclei; area under the receiver-operating-characteristic curve [AUC] = 0.51; Fig. 3l). In the corresponding super-resolution images, malignant nuclei exhibited lower homogeneity and substantially greater separation from benign nuclei (Cliff’s δ = 0.46; AUC = 0.73; Fig. 3m). Other GLCM texture features, such as contrast, showed a similar improvement in separability under super-resolution, whereas intensity-and morphology-based features remained weakly discriminative under both imaging conditions (**Supplementary Fig. S6**). Thus, the improvement was concentrated in local nuclear texture rather than global nuclear intensity or morphology. These results indicate that MOSH recovered nuclear-texture information that was blurred in the corresponding wide-field images and demonstrate its potential to enhance quantitative nuclear phenotyping(*51–53*).

Finally, the reconstructed survey image served as a spatial map for targeted three-dimensional interrogation of selected benign and malignant regions. MOSH returned to the mapped specimen locations and acquired super-resolution volumes, revealing the depth-dependent arrangement, clustering, and morphological heterogeneity of nuclei within their local tissue environments (**Fig. 3n-q**). Targeted volumetric imaging resolved the three-dimensional nuclear architecture within histologically defined benign (**Fig. 3n, o**) and malignant (**Fig. 3p, q**) regions, revealing heterogeneity in nuclear position and morphology throughout the local tissue depth. Sequential optical sections further resolved nuclei located at different axial depths that were simultaneously represented in the depth-extended survey image, demonstrating depth-resolved structural separation within the tissue (**Supplementary Fig. S7**). Across both tissue morphologies, wide-field volumes exhibited broad voxel-intensity distributions due to out-of-focus background. MOSH shifted these distributions toward lower background intensities while retaining structural signal, consistent with improved optical sectioning in dense, scattering tissue (**Fig. 3r, s; Methods, Supplementary Note 4**). Collectively, these results demonstrate that MOSH connects millimeter-scale histological context, quantitative subnuclear texture, and targeted three-dimensional architecture within a single spatially registered workflow.

### Section-free super-resolution optical biopsy of intact tissue

Conventional histopathology relies on physical sectioning to expose internal tissue architecture, providing a standardized diagnostic workflow at the expense of preserving the intact three-dimensional specimen(*14, 54*). Here, we evaluated MOSH using mouse kidney tissue prepared as conventional formalin-fixed, paraffin-embedded (FFPE) sections and as unsectioned punch biopsies from perfused and non-perfused kidneys (**Fig. 4a, Methods**). MOSH produced large-area super-resolution fluorescence and corresponding virtual H&E images of FFPE sections, based on compatible, established histological preparations (**Fig. 4b-g, Supplementary Fig. S8**). In contrast to the multistep processing required for FFPE tissue, the intact biopsies were labeled directly with DRAQ5 and Eosin Y in less than 10 min and imaged without physical sectioning, thereby preserving the tissue architecture for subsequent optical interrogation (**Methods, Supplementary Note 5**).

**Fig. 4.**
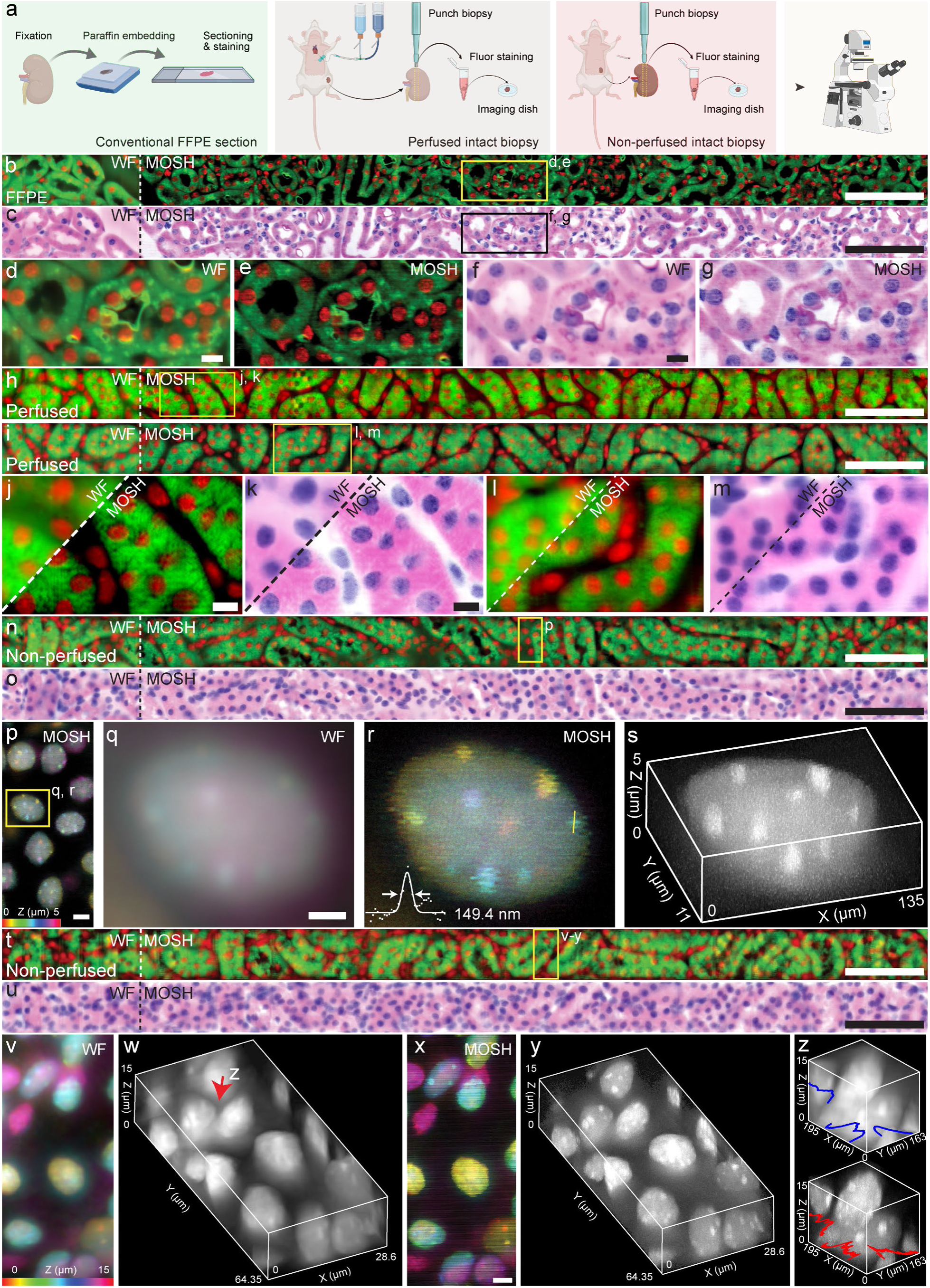
Sectioned and section-free MOSH imaging of the mouse kidney. (a) Schematic of the preparation workflows for conventional formalin-fixed, paraffin-embedded (FFPE) sections and intact punch biopsies obtained from perfused and non-perfused mouse kidneys. Red, DRAQ5; green, Eosin Y. (b, c) 5-µm extended-depth-of-focus (EDoF) survey of an FFPE kidney section, shown as fluorescence (b) and corresponding virtual H&E (c) images. (d–g) Zoomed-in wide-field (WF) and MOSH images of the boxed regions in (b, c), shown as fluorescence (d, e) and virtual H&E (f, g) images. (h–m) EDoF surveys of intact perfused biopsies across tunable axial ranges of 2.25 µm (h) and 5 µm (i), with diagonally divided zoomed-in WF and MOSH images shown as fluorescence (j, l) and virtual H&E (k, m) images. (n, o) 5-µm EDoF survey of an intact non-perfused biopsy containing residual blood, shown as fluorescence (n) and corresponding virtual H&E (o) images. (p) Depth-color-coded MOSH projection of the nuclear channel from the boxed region in (n), with color denoting axial position. (q, r) WF (q) and MOSH (r) views of the boxed nucleus in (p). The line profile in (r) indicates an apparent lateral width of approximately 150 nm for the selected intranuclear feature. (s) MOSH volumetric reconstruction of the nucleus selected in (p). (t, u) 15-µm EDoF survey of intact non-perfused kidney tissue, shown as fluorescence (t) and corresponding virtual H&E (u) images. (v–y) Depth-color-coded projections (v, x) and volumetric reconstructions (w, y) of the boxed region in (t) under WF (v, w) and MOSH (x, y) imaging. (z) WF (top) and MOSH (bottom) volumetric comparison of the nucleus indicated by the arrow in (w), with line-intensity profiles along the indicated paths. Scale bars: 100 µm (b, c, h, i, n, o, t, u), 10 µm (d–g, j–m), 5 µm (p, v, x), and 2 µm (q, r).

In perfused biopsies, we demonstrated that MOSH generated super-resolved survey images across tunable axial ranges (**Fig. 4h, i**), revealing nuclear and extracellular matrix structures with greater clarity than the corresponding wide-field images (**Fig. 4j-m**). Sequential optical sections resolved the depth-dependent distribution of individual nuclei, complementing the depth-integrated survey images with spatially resolved axial information (**Supplementary Fig. S8**). We then applied the same workflow to non-perfused biopsies, in which residual blood introduced additional absorption, scattering, and fluorescence background. MOSH preserved nuclear and renal microstructural contrast across a 5-µm axial range of renal tissue (**Fig. 4n**), and virtual H&E rendering transferred the recovered subcellular information into a commonly adopted histological representation (**Fig. 4o**). We next used the reconstructed survey image as a spatial map for targeted volumetric interrogation of selected nuclei (**Fig. 4p**). Within a survey-identified region, targeted volumetric imaging resolved the three-dimensional organization of nuclei and revealed DRAQ5-negative intranuclear features, morphologically consistent with nucleoli, which were obscured by wide-field imaging (**Fig. 4q, r**). The results resolved nuclear features at approximately 150 nm and localized them to distinct axial positions throughout the nucleus, demonstrating the recovery of subdiffractional structural information within intact tissue (**Fig. 4s, Supplementary Note 5**).

Finally, we extended MOSH to image an axial range of 15 µm within unsectioned, non-perfused kidney tissue containing residual blood (**Fig. 4t, u**). Under these extensive conditions, targeted super-resolution reconstruction preserved optical sectioning and resolved nuclear boundaries and intranuclear structures, exhibiting lower background and greater spatial modulation than wide-field imaging (**Fig. 4v-z**). Collectively, these results establish MOSH as a unified framework that extends super-resolution histology from conventional tissue sections to rapid, section-free optical biopsy, integrating large-area screening, virtual H&E visualization, and targeted volumetric interrogation of intact tissue.

## DISCUSSION

Super-resolution microscopy reveals subcellular tissue architecture obscured by conventional imaging, yet extending this capability to histological screening requires balancing spatial resolution with imaging area, axial access, throughput, and instrumental complexity(*55, 56*). Resolving these challenges remains essential for translating super-resolution imaging from a specialized microscopy modality into a practical tissue assessment tool. In this work, MOSH integrates depth-extended super-resolution survey imaging with coordinate-guided three-dimensional interrogation in conventional epi-fluorescence settings. Its survey-and-target strategy preserves tissue-scale context while concentrating volumetric acquisition on selected subcellular structures. Automated reconstruction and virtual H&E rendering connect multiscale acquisition with a familiar histological representation. Further advances in optical design(*57–59*), learning-based reconstruction(*60–63*), and specimen handling(*64–68*) could improve imaging performance, computational latency, and specimen compatibility. Integrating MOSH with complementary microscopy architectures(*69–71*), spatial omics(*72, 73*), and computational pathology(*74, 75*) could enable adaptive, structure-guided workflows in which survey-scale morphology directs subsequent molecular and volumetric interrogation. We anticipate that MOSH will provide a scalable bridge between tissue-scale histological screening and quantitative nanoscale interrogation across biological and clinical specimens.

## MATERIAL AND METHODS

### Imaging setup

In this study, fluorescence was excited using 647-nm and 488-nm lasers, with the beam expanded and uniformly shaped by a pinhole positioned between 30-mm and 500-mm achromatic lenses. This collimated beam passed through a microlens array (MLA) to generate a grid of multifocal spots, which were directed by a dichroic mirror into an electro-tunable lens (ETL). An additional 1:1 relay pair then positioned the beam precisely at the conjugate back focal plane of the objective, where it passed through the microscope’s backport relay arm to generate a grid of diffraction-limited spots on the sample. The emitted fluorescence was collected by the objective and returned along the same path, at which point the system used one of two distinct acquisition methods. For 2D extended depth-of-field (EDoF) imaging, the ETL actively scanned the excitation beam and de-scanned the returning emission before it was separated by the dichroic mirror, focused by a tube lens, and detected by an sCMOS camera. Alternatively, for volumetric acquisition, the ETL was held static at its equilibrium position, and a piezo Z-stage was instead used to sequentially step the objective lens along the Z-axis to capture discrete depth slices. A complete list of optical components and hardware is provided in **Supplementary Table S2.**

### High-speed image acquisition pipeline

An automated Python pipeline was developed to control image acquisition and reconstruction using Micro-Manager through the Pycro-Manager library. Hardware-level external triggering between the XY stage and camera was used to synchronize stage motion and image acquisition. Concurrent acquisition and data writing were implemented using a multithreaded producer–consumer architecture to maintain continuous data transfer during high-throughput imaging.

An 8-GB circular buffer was allocated within the Micro-Manager core to accommodate transient fluctuations in disk-write speed and reduce the likelihood of dropped frames. Acquired NumPy arrays were stored using the Zarr format rather than as individual large monolithic files. A dedicated background writer thread continuously retrieved frames from the Micro-Manager buffer, organized them into 128-frame chunks, and wrote them to disk using Blosc LZ4 compression while acquisition continued. Following acquisition, the Zarr datasets were rechunked using Dask from 128-frame to 300-frame blocks to match the block size used for downstream reconstruction and improve reconstruction I/O efficiency.

The acquisition sequence was performed independently for the two fluorescence channels. The 488-nm channel was acquired first while background data writing proceeded concurrently. Upon completion, the stage returned to the acquisition origin and restored the designated scanning speed before acquisition of the 647-nm channel. The resulting channel-specific Zarr datasets were subsequently passed to the reconstruction pipeline together with their corresponding mapping files.

### Image reconstruction

The reconstruction process begins by evaluating the raw data to verify that the stage’s frame-by-frame translation across the sample was uniform. To mitigate motion-induced artifacts caused by mechanical stage deviations, a two-stage digital tracking protocol was applied to the raw image stacks. First, tracking along the scanning direction was performed using zero-mean normalized cross-correlation (ZNCC) template matching. To account for non-uniform sample illumination and continuous downward shifting of the sample, a dynamic region of interest (ROI) was implemented. By progressively lowering the upper boundary of the search window frame-by-frame, the search region height was dynamically restricted, significantly reducing false matches in irrelevant regions. When additional correction for drift in the non-scanning axis was required, Enhanced Correlation Coefficient (ECC) registration was applied to the 1D-aligned image stack. The mean of the aligned frames served as the registration template, and ECC iteratively estimated a translation-only transformation for each frame to correct residual 2D drift.

Following image registration, an illumination map (foci map) was applied to isolate individual emission spots. Second, digital pixel reassignment was performed by reassigning the shifted pixels to the theoretical pinhole positions, yielding an intermediate image with a theoretical 1.4× resolution improvement. Finally, a rapid, single-iteration Wiener-Butterworth deconvolution was applied using a theoretically derived point spread function (PSF) to achieve the full 2× resolution enhancement (**Supplementary Note6, Supplementary Fig. S9**). In tissue samples exhibiting highly non-uniform illumination, pixel reassignment occasionally induced high-frequency striping artifacts. Where necessary, a post-processing correction was implemented by extracting a normalized median projection of the image stack, applying a high-pass filter to isolate the striping pattern, and performing flat-field division using this isolated artifact mask.

### High-throughput software architecture

To process datasets larger than the available system memory, the reconstruction pipeline was implemented using a controlled nested-parallelism architecture with Dask. At the data-loading level, Zarr metadata were used to identify the required chunks, which were loaded sequentially as needed rather than loading the complete dataset into memory. At the processing level, reconstructed substacks were distributed across a local worker cluster for parallel computation.

For quality control, the first data substack was reconstructed independently using a pre-calibrated fixed scan-angle parameter, providing a rapid preview without performing full 1D trajectory tracking. If the preview showed adequate tracking, the fixed-angle parameter was retained for subsequent reconstruction. If tracking deviations were observed, full 1D ZNCC tracking was performed to estimate the scan trajectory, with ECC-based 2D translational refinement applied when additional correction was required. Once the tracking approach was established, the remaining frames were partitioned into overlapping substacks with a one-frame overlap and distributed across the local worker cluster. This combination of substack-level parallel processing and sequential disk I/O enabled high-throughput reconstruction while limiting system-memory requirements.

### Morphology-guided target prioritization and coordinate transformation

Following 2D survey reconstruction, individual nuclei within each reconstructed survey tile were segmented using the Cellpose nuclei model. For the targeting implementation used here, nuclear area and eccentricity were extracted from each segmentation and standardized within each tile using Z-scores. The standardized values were summed to generate a morphometric prioritization score, with higher scores corresponding to nuclei that were larger and more eccentric relative to the local nuclear population. Each survey tile was ranked according to its highest-scoring nucleus, and the three highest-ranked tiles were presented to the user through a custom graphical interface.

The interface displayed the reconstructed image together with the Cellpose segmentation and highlighted the highest-scoring candidate nuclei to guide review. The user then manually delineated a region of interest (ROI) within the selected tile for subsequent 3D imaging. This analysis step was implemented as one example of morphology-guided target prioritization; alternative image-derived features or scoring strategies can be incorporated into the same targeting workflow.

The selected image coordinates were converted to absolute lateral stage coordinates using a calibrated coordinate transformation that accounted for the physical origin of the corresponding survey tile, the camera ROI offset, and the effective pixel size of the reconstructed SR image. The resulting (X, Y) coordinates were then passed to the targeted volumetric acquisition pipeline.

### High-speed targeted volumetric acquisition

For targeted volumetric imaging, the automated pipeline directed the translation stage to the absolute coordinates of the user-selected ROIs. The camera was operated in external hardware-trigger mode, with exposures initiated by TTL pulses from the stage controller to synchronize image acquisition with stage motion.

Upon arrival at each target coordinate, a temporary local coordinate origin was established in the stage controller, enabling relative scanning over the selected subregion. The stage then performed a continuous lateral scan at successive axial (Z) planes while generating hardware triggers for synchronized camera acquisition. Following completion of the volumetric scan, the global coordinate system was restored, and the stage advanced to the next selected ROI, where the acquisition sequence was repeated.

### Virtual hematoxylin and eosin (H&E) staining

Following reconstruction, the high-resolution fluorescence substacks were merged into large-field-of-view mosaics using the Grid/Collection Stitching plugin in Fiji/ImageJ(*76*). Dual-channel fluorescence images were then converted to virtual hematoxylin and eosin (vH&E) images using a Beer–Lambert-based rendering algorithm. The nuclear and cytoplasmic fluorescence channels were used as spatial concentration maps for hematoxylin-like and eosin-like staining, respectively.

Each channel was normalized to its 99.9th-percentile intensity to reduce the influence of isolated high-intensity pixels. For the nuclear channel, normalized values below 10% were set to zero, and the remaining intensities were rescaled to suppress low-level background. Nonlinear gamma correction was then applied using fixed exponents of 1.5 for the nuclear channel and 1.2 for the cytoplasmic channel.

The processed concentration maps were converted to optical density (OD) using hematoxylin and eosin color vectors from Ruifrok and Johnston (*77*). Nuclear and cytoplasmic OD contributions were weighted by factors of 1.9 and 1.0, respectively, and combined before conversion to RGB intensity using the Beer–Lambert relationship.

### Quantitative feature analysis and group separation

Group separation was quantified using Cliff’s delta (δ) and the area under the receiver operating characteristic curve (AUC). Cliff’s δ ranges from −1 to +1, where 0 indicates complete distributional overlap and larger absolute values indicate greater separation between groups. Using nuclei sampled from benign tissue regions as the reference group, positive δ values denote lower feature values in nuclei sampled from malignant regions. AUC values range from 0.5, indicating chance-level rank separation, to 1.0, indicating complete separation. When imaging modalities were compared, these separability metrics were computed independently within each modality.

Volumetric background rejection was assessed by comparing voxel-intensity distributions obtained from three-dimensional nuclear regions of interest in wide-field and MOSH volumes. Details of the volumetric intensity analysis are provided in **Supplementary Note 4**.

### Suspension cell line (K562) preparation

Cryopreserved K562 cells were generously provided by Dr. Gabe Kwong’s laboratory and thawed according to standard recovery protocols prior to seeding. Cells were maintained in Iscove’s Modified Dulbecco’s Medium (IMDM) supplemented with 10% fetal bovine serum (FBS) at 37 °C and 5% CO₂ and routinely passaged during culture. K562 cells were prepared using three cytological preparation formats: glass-bottom imaging dishes, coverslips mounted onto glass slides, and direct slide smears. For adherent preparations, glass-bottom imaging dishes (µ-Dish) were coated with 1 mL of 0.01% poly-L-lysine for 5 min at room temperature. Following aspiration and a 30-min desiccation period, K562 cells were seeded in a 1 mL volume and incubated for 30 min. Cells were fixed in 4% paraformaldehyde (PFA) for 10–15 min, washed three times in 1× PBS, and incubated in 5–10 µM DRAQ5 for 10 min at room temperature. After a 1× PBS wash, cells were stained with 0.5% (w/v) Eosin Y for 30 s, washed three to four times in 1× PBS, and maintained in 1× PBS for imaging.

For slide and coverslip preparations, K562 cultures were centrifuged at 300 × g for 5 min. The supernatant was decanted to yield an approximate residual volume of 50 µL for cell resuspension. Suspensions were applied directly to coverslips or mechanically smeared onto glass slides, air-dried for 1 min, and fixed in either absolute methanol for 1 min or 4% PFA for 10 min. Following a gentle 1× PBS wash, cells were incubated with 50 µM DRAQ5 for 10 min. The stain was decanted, and the preparation was rinsed with 1× PBS prior to incubation with Eosin Y for 30–60 s. Slides were cleared in distilled water or 1× PBS. Coverslip preparations were mounted onto glass slides using a tape spacer to prevent mechanical compression of the cells.

### Standard FFPE tissue staining

De-identified, archived formalin-fixed, paraffin-embedded (FFPE) human tissue sections were obtained from the Cancer Tissue and Pathology Shared Resource of Winship Cancer Institute of Emory University. The use of these fully de-identified retrospective specimens was reviewed by the Emory University Institutional Review Board (IRB) and determined to be Non-Human Subjects Research; therefore, formal IRB approval and informed consent were not required. The samples utilized consisted of two colon adenocarcinoma (not otherwise characterized) specimens: a Stage I (pT2N0) specimen from a 67.7-year-old White, non-Hispanic male, and a Stage II (pT3N0) specimen from a 52.9-year-old Black or African American, non-Hispanic male.

FFPE tissues were sectioned at 5 µm, mounted onto glass slides, and deparaffinized and rehydrated according to standard protocols. For nuclear labeling, slides were incubated in 50 µM DRAQ5 in 1× PBS for 7 min at room temperature, followed by a 1-min wash in 1× PBS. Sections were subsequently counterstained in 0.2–0.5% (w/v) Eosin Y in 95% ethanol for 15–20 s with continuous agitation. Differentiation was performed in 95% ethanol for 30–60 s until background signal on the glass substrate was eliminated. Slides were dehydrated in absolute ethanol for 10 s, excess solvent was aspirated from the slide periphery, and sections were mounted in 100% glycerol and sealed.

### Mouse kidney FFPE processing

All animal husbandry and experimental procedures were performed in accordance with protocol PROTO202200102 approved by the Emory University School of Medicine Institutional Animal Care and Use Committee (IACUC). Mouse kidney tissues were obtained from Dr. Alan Emanuel’s laboratory via postmortem sharing of animal tissue. Tissues were collected postmortem from animals that had been humanely euthanized for independent experimental purposes. Extracted kidneys were post-fixed in 10% neutral buffered formalin, dehydrated through a graded ethanol series, cleared in xylene, and embedded in paraffin. Tissues were sectioned at 5 µm using a rotary microtome (Epredia HM 340E Microtome B) and mounted onto glass slides. Deparaffinization and rehydration were performed using an Autostainer XL (Leica Biosystems), consisting of xylene clearance (two changes, 10 min each), rehydration through absolute ethanol (two changes, 10 min each), followed by 95%, 70%, and 50% ethanol (5 min each), and a final 10-min equilibration in 1× PBS. Sections were subjected to a sequential dual-staining protocol: a 10-s incubation in 50 µM DRAQ5, a 1× PBS rinse, and a 10-s counterstain in 2% (w/v) Eosin Y in 80% ethanol. Following a brief 1× PBS wash, a secondary 3-min incubation in 50 µM DRAQ5 was performed to amplify the nuclear signal. Slides were rapidly dehydrated through a graded ethanol series (70%, 95%, 100%), cleared in xylene, and mounted with a fluorescence-compatible mounting medium.

### Fresh mouse kidney biopsies

Fresh mouse kidney tissues were similarly obtained from Dr. Alan Emanuel’s laboratory through postmortem sharing of animal tissue, in accordance with the Emory University IACUC protocol (PROTO202200102). For perfused tissue models, mice were subjected to deep surgical anesthesia with isoflurane followed by cervical dislocation, and systemic perfusion was performed via the left ventricle using cold 1× PBS to achieve complete exsanguination prior to organ harvest. Unperfused control models were harvested immediately following isoflurane anesthesia and cervical dislocation. Extracted organs were submerged in cold 1× PBS, placed in an ice container, and transported directly to the laboratory.

Following transport, 4-mm tissue cores were extracted using a dermal punch biopsy (Model MB, MedBlades) for immediate whole-mount fluorescence imaging. Both perfused and unperfused tissues were briefly washed in 1× PBS; unperfused samples received multiple additional cold 1× PBS rinses to clear excess luminal and surface blood prior to staining. Biopsies were then submerged in Eosin Y (0.1–0.2% w/v) for 30 seconds to stain the cytoplasm and extracellular matrix, followed by two brief rinses in 1× PBS. Samples were next incubated in a 10–20 µM DRAQ5 solution for 10–15 minutes at room temperature to selectively label cell nuclei, followed by a final 1-minute 1× PBS wash to clear background fluorescence. To preserve 3D architecture while achieving a flat imaging plane, the stained biopsies were mounted in a glass-bottom dish using a capillary-lock technique, in which a 50–100 µL droplet of PBS and a square coverslip were applied directly over the tissue to create a level focal surface under surface tension. To minimize tissue degradation, all transfer, staining, and initial imaging procedures were completed within a 2-hour window following organ harvest. Mouse kidney tissues were handled under an approved Georgia Institute of Technology Biological Materials Safeguards Committee (BMSC) protocol (Permit No. B-937-B-001).

## Acknowledgments

We thank Alan Emanuel and his laboratory for providing the mouse kidney tissue samples used in this study. We also thank the Georgia Tech Histology Core for assistance with tissue processing.

## Author contributions

Conceptualization: K.T., S.J. Methodology: K.T. Software: K.T., O.L.

Investigation: K.T., Q.L., V.P., R.Z.

Formal analysis: K.T., H.Y., E.C. Validation: K.T., E.C.

Resources: B.R., J.R., P.C., R.Z., Z.Z., G.K.

Visualization: K.T., P.C. Supervision: S.J. Writing—original draft: K.T., S.J.

Writing—review & editing: K.T., H.Y., S.J.

## Data and materials availability

All data needed to evaluate the conclusions in the paper are present in the paper and/or available upon request from the corresponding author. MOSH is available as Supplementary Software. The code is written in Python 3.11.4. The latest version of the software is available at https://github.com/ShuJiaLab/MOSH.

