## Supplemental information for "Multiscale, Survey-and-Target Optical Super-Resolution Histology"

Kidan Tadesse *et al.*

**This PDF file includes:**

Supplementary Notes Sections S1 to S6

Figs. S1 to S9

Tables S1 to S3

References (1-10)

SUPPLEMENTARY NOTE

**Note S1**

MOSH System Architecture and Acquisition Performance

To enable high-throughput multiscale imaging, the MOSH optical setup (**Fig. S1**) is governed by a synchronized, four-zone computational architecture **(Fig. S2)**. This centralized platform orchestrates the transition between hardware states, shifting from active electrically tunable lens (ETL) sweeping for macroscopic screening to active Z-stage translation for targeted volumetric capture (**Fig. S2A**). To manage the large data volumes generated during the initial screening phase, the software utilizes a multi-threaded Producer-Consumer architecture to execute simultaneous hardware-level TTL triggering, data compression, and disk writing (**Fig. S2B, Zone 1**). In a representative macroscopic experiment, this pipeline enabled the imaging of a 1,179 × 65 µm tissue strip in ~100 s at a 100 Hz camera framerate. During continuous acquisition, the asynchronous data-writing framework compressed a ~38 GB raw dataset into memory-safe Zarr arrays of ~10 GB, eliminating temporal overhead and preventing throttling of the optical capture rate.

Following acquisition, the pipeline employs a distributed Dask framework to parallelize the super-resolution reconstruction (**Fig. S2B, Zone 2**). This deterministic workflow (**Materials and** **Methods**) yields a twofold improvement in lateral resolution, resolving subcellular features down to 120–150 nm across an extended depth of field (EDoF) of up to 20 µm. By replacing iterative deconvolution protocols with a single-pass filter, the pipeline reconstructed the complete macroscopic field of view in ~6 min per fluorescence channel. The reconstructed data directly feed the subsequent AI-driven morphological targeting loop and virtual H&E visualization pipeline (**Fig. S2B, Zones 3 and 4**), providing the computational efficiency necessary for routine multiscale tissue evaluation.

.

**Note S2**

Quantitative Analysis and Additional Cytological and Hematological Imaging

MOSH achieved a lateral survey throughput of 766 µm² s⁻¹. Because EDoF acquisition is performed simultaneously with lateral scanning, extending the EDoF range up to 20 µm does not increase acquisition time, corresponding to a maximum depth-extended survey throughput of 15,320 µm³ s⁻¹. A 5 µm EDoF was used for the cytological preparations in Fig. 2 to match the required specimen depth rather than an acquisition-speed limitation.

Lateral image resolution in the extended-depth-of-field (EDoF) cytology images was estimated using decorrelation analysis (DeARes)(*1*) of individual K562 cells from each preparation format. Ten cells were analyzed independently for each condition. The resulting mean lateral resolution estimates were 140.3 nm for cells cultured in a dish, 140.6 nm for cells mounted on a slide, and 199.5 nm for cells prepared as a smear (n = 10 cells per measurement).

To quantify nuclear structural definition, individual nuclei were identified by automated thresholding and watershed segmentation. The mean nuclear intensity gradient was then calculated from Sobel-filtered images to quantify local image sharpness within each segmented nucleus. Nuclear gradients were compared between MOSH super-resolution and conventional wide-field images across the dish, slide, and smear preparation formats. MOSH produced significantly higher mean nuclear intensity gradients across all three preparation formats (P < 0.001, Welch’s t-test), with the largest difference observed for the dish and smear preparations.

Building upon the volumetric acquisition of intact K562 cells in a dish, additional 3D reconstructions were performed to visualize isolated nuclear topologies (**Fig. S3**). A depth-color-coded projection of a K562 nucleus (**Fig. S3A**) and the corresponding volumetric reconstruction (**Fig. S3B**) visualize its three-dimensional surface morphology. Sequential super-resolution optical sections acquired across a 5.25 µm depth range (**Fig. S3C**) further reveal depth-dependent changes in nuclear morphology.

To further evaluate continuous EDoF imaging in a cell-dense cytological preparation, MOSH was applied to a standard blood smear using an eosin-only contrast channel (**Fig. S3D**). A continuous field of view of 1157.73 × 118.95 µm was acquired with a 5 µm EDoF in approximately 200 s, corresponding to an area throughput of approximately 690 µm² s⁻¹. Subsequent Cellpose segmentation identified 560 individual cells within the acquired strip, corresponding to an observed survey rate of approximately 2.8 cells s⁻¹. These results demonstrate large-area EDoF acquisition with downstream automated cell segmentation in a densely populated hematological preparation.

**Note S3**

Spatial Fidelity and Optical Sectioning in Clinical Histology

To further characterize MOSH imaging in clinical tissue, large-field-of-view imaging was performed on a stage II human colon cancer section over a 5 µm extended depth of field (EDoF) (**Fig. S5A, B**). Line-intensity profiles were extracted from the native fluorescence images and corresponding virtual H&E representations (**Fig. S5C-F**) to evaluate preservation of spatial features during computational translation. Cross-sectional profiles from the wide-field (**Fig. S5I**) and MOSH (**Fig. S5J**) images showed spatial correspondence between features in the native fluorescence and virtual H&E images, indicating that the virtual staining procedure retained the spatial organization of the underlying fluorescence data.

Short-range intensity profiles were further used to compare the separation of closely spaced structural features (**Fig. S5G, H**). Features that appeared as a broad unresolved intensity distribution under wide-field imaging showed increased modulation and distinct peak separation in the corresponding MOSH profiles. Similar differences were retained after virtual H&E translation, supporting preservation of the additional spatial information recovered in the super-resolution fluorescence images.

Optical sectioning was evaluated over an approximately 30 µm profile spanning densely packed nuclei (**Fig. S5M, N**). Wide-field profiles exhibited an elevated background and reduced modulation between adjacent structures, whereas the corresponding MOSH profiles showed lower background levels and deeper intensity minima between neighboring features. These differences are consistent with reduced contribution from out-of-focus fluorescence in the MOSH images.

Image resolution was further estimated using decorrelation analysis2. Within the multiplexed tissue region shown in (**Fig. S4E)**, the combined image yielded an estimated structural resolution of approximately 212.7 nm. Channel-specific analysis yielded estimates of approximately 182.7 nm for the nuclear channel and 178.9 nm for the cytoplasmic channel. In the isolated nuclear region shown in (**Fig. S5K)**, decorrelation analysis yielded an estimated resolution of 209 nm for MOSH compared with 669 nm for the corresponding wide-field image. Together, these measurements support improved spatial resolution and reduced out-of-focus background in MOSH imaging of clinical colon tissue.

**Note S4**

Colon tissue nuclear texture analysis

Nuclei were segmented using Cellpose (version 4.1.13) with a diameter of 80 pixels, flow threshold of 0.4, and cell probability threshold of 0. Identical segmentation parameters were applied across all four images. Objects smaller than 300 pixels or larger than the 99.7th percentile of the area distribution were excluded, yielding 927 benign and 569 malignant nuclei in WF and 942 benign and 543 malignant nuclei in SR.

Feature extraction was restricted to pixels within each nuclear mask. For texture analysis, masked gray-level co-occurrence matrices (GLCMs) were constructed using only pixel pairs for which both pixels were located within the same nuclear mask. Intensities were quantized to 32 gray levels, and GLCMs were calculated at a displacement of 1 pixel in four directions. Homogeneity, contrast, and correlation were extracted from the resulting GLCMs. Additional features included mean intensity, intensity standard deviation, intensity entropy, nuclear area, eccentricity, and aspect ratio.

Benign–malignant group separation was quantified independently within each imaging modality using Cliff’s delta (δ) and the area under the receiver operating characteristic curve (AUC). Cliff’s δ ranges from −1 to +1, with values near zero indicating substantial overlap and larger absolute values indicating greater separation. Using benign nuclei as the reference group, positive δ values denote lower feature values in malignant nuclei. AUC values of 0.5 and 1.0 indicate chance-level and complete rank separation, respectively. For nuclear homogeneity, WF imaging showed essentially no benign–malignant separation (δ = −0.02, AUC = 0.51), whereas SR imaging produced substantially greater separation (δ = 0.46, AUC = 0.73), with malignant nuclei exhibiting lower homogeneity. Contrast and correlation showed similar modality-dependent increases in group separation, whereas intensity- and morphology-based features remained weakly discriminative. Full feature distributions are provided in **Fig. S6.**

To evaluate volumetric background rejection and optical sectioning between wide-field (WF) and MOSH imaging, three-dimensional regions of interest (ROIs) tightly bounding individual nuclei were cropped from the acquired volumes. For each volumetric ROI, the intensities of all voxels were pooled to generate a voxel-intensity distribution in MATLAB. The resulting distributions were compared between imaging modalities to assess the relative contributions of background and structural signal. WF volumes exhibited broad intensity distributions displaced from zero, consistent with contributions from out-of-focus fluorescence throughout the volumetric ROI. In comparison, MOSH volumes showed an increased population of low-intensity background voxels while retaining a higher-intensity component associated with nuclear structure, consistent with improved volumetric background rejection and optical sectioning.

**Note S5**

Volumetric imaging and optical sectioning of intact mouse kidney tissue

To further evaluate the relationship between depth-extended survey imaging and volumetric acquisition, additional analysis was performed on the FFPE mouse kidney specimen shown in Fig. 4B (**Fig. S8A-D**). The MOSH EDoF image projects fluorescence collected over the acquired axial range into a single depth-extended representation, allowing nuclei located at different axial positions to remain visible within the survey image (**Fig. S8A**). Targeted volumetric acquisition of a selected region was subsequently visualized as a depth-color-coded projection and three-dimensional reconstruction (**Fig. S8B, C**). Examination of sequential optical sections showed individual nuclei entering and leaving the focal plane at different axial positions (**Fig. S8D**), demonstrating recovery of depth-dependent structural information that is not retained in the EDoF representation alone.

Additional volumetric analysis was performed on intact perfused mouse kidney punch biopsies acquired over axial ranges extending 2.25 and 5 µm from the tissue surface (**Fig. S8E-L**). At 2.25 µm, depth-color-coded projections showed reduced out-of-focus background in the MOSH reconstruction relative to wide-field imaging, while the corresponding volume rendering visualized the three-dimensional organization of nuclei within the selected region (**Fig. S8E-H**). Similar differences were retained over the 5 µm imaging depth (**Fig. S8I-K**). Sequential super-resolution optical sections through selected nuclei further revealed depth-dependent changes in nuclear morphology and separation of structures located at different axial positions (**Fig. S8L**).

Zoomed-in analysis of the targeted nucleus shown in Fig. 4Q, R revealed intranuclear features consistent with nucleoli that were not distinctly resolved in the corresponding wide-field image. To quantify the lateral dimension of the selected feature, a line-intensity profile was extracted across the structure. Analysis of the resulting profile yielded an apparent lateral feature dimension of 149.38 nm. This value represents the measured dimension of the selected biological structure rather than an independent estimate of the intrinsic system resolution.

**Text S6**

Validation of Non-Iterative Super-Resolution Reconstruction

To accommodate the massive data throughput of the MOSH platform, the computational pipeline must prioritize deterministic, single-pass algorithms over traditional iterative approaches. While iterative blind deconvolution remains a conventional method for resolving sub-diffraction structures, it imposes a severe computational bottleneck when scaled to the gigapixel volumetric datasets required for clinical histology.

To validate our streamlined computational approach, we evaluated a rapid, single-iteration Wiener-Butterworth (WB) deconvolution(*2*) against a previously utilized iterative blind deconvolution method using a standardized microtubule MOSH dataset (**Fig. S9**). Image decorrelation analysis quantitatively confirmed the robust performance and efficiency of the WB filter. While the diffraction-limited widefield baseline measured at ~410 nm, both deconvolution methods successfully pushed the resolving power deep into the sub-diffraction regime. Notably, the WB filter yielded a lateral resolution of ~177 nm, matching and slightly outperforming the ~187 nm resolution achieved by the iterative blind method, while preserving equivalent image contrast and structural delineation. Crucially, whereas blind deconvolution required approximately 5 iterations to reach this image quality, the deterministic WB filter achieved it in a single pass. This corresponds to a ≥5× improvement in deconvolution speed with no penalty to the final spatial resolution, demonstrating the absolute suitability of the WB architecture for the high-throughput, memory-safe reconstruction of massive tissue volumes.

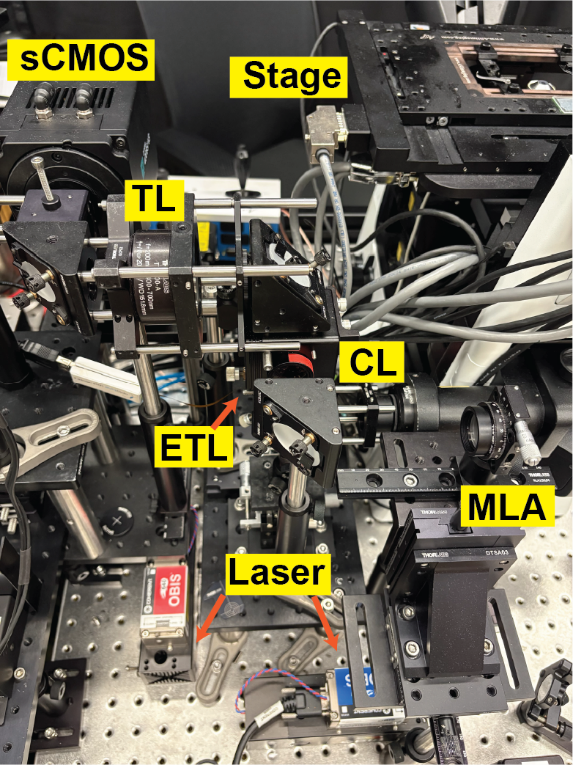

Fig. S1. MOSH experimental setup. The light beam (488-nm and 647-nm lasers) is expanded, shaped by a pinhole, and passed through the microlens array (MLA) that generates multifocal excitation patterns. The beam is directed by a dichroic mirror through the electrically tunable lens (ETL) and relayed to the objective lens. The excited sample is continuously translated by the motorized X-Y stage. For 2D extended depth of field (EDoF) imaging, the sample is scanned in the Z direction by the ETL; the emitted light from the sample is collected by the objective lens, de-scanned by the ETL, and recorded by the sCMOS camera. For targeted volumetric acquisition, the ETL is held static, and the sample is instead scanned in the Z direction by a piezoelectric Z-stage to capture discrete depth slices.

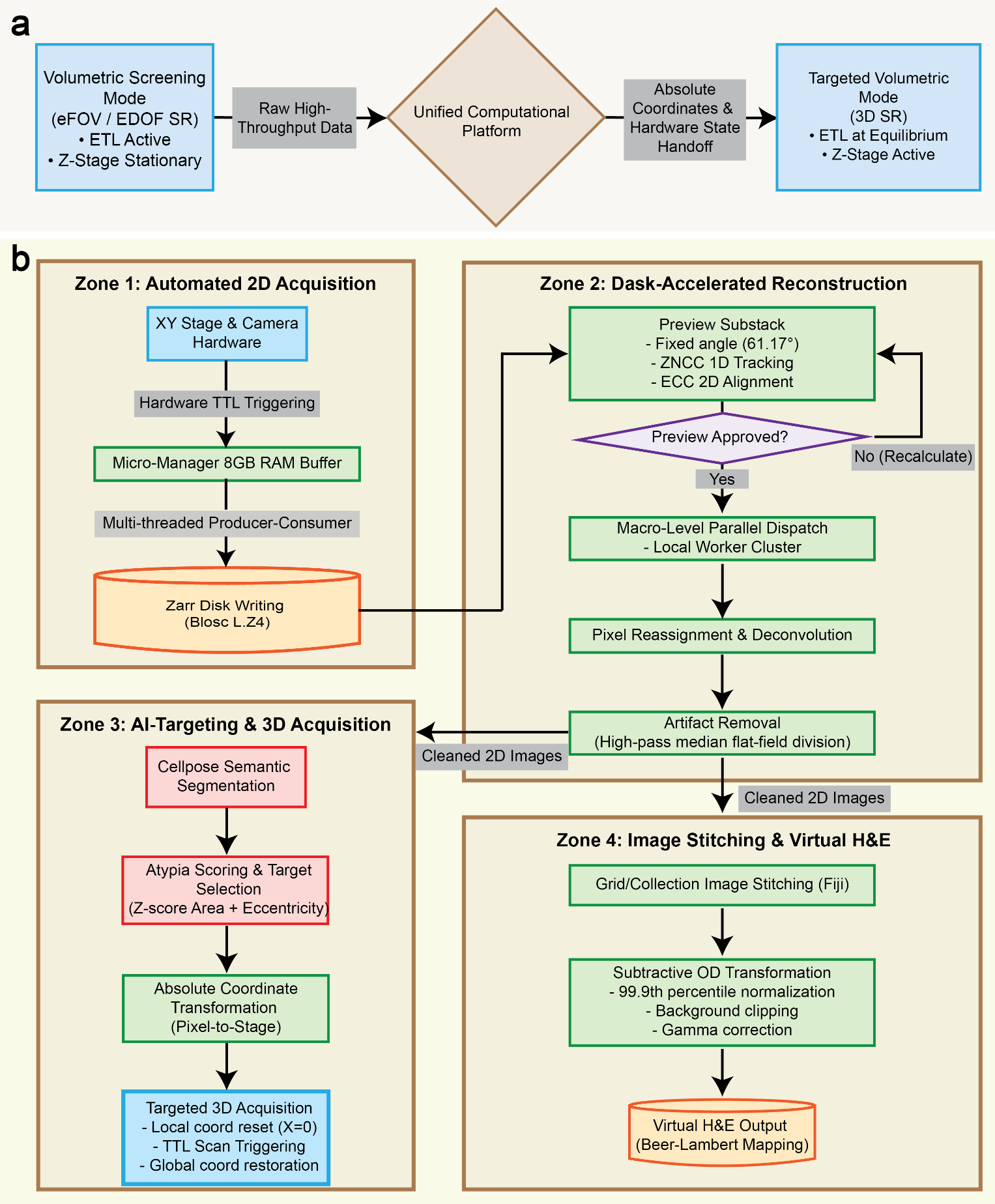

Fig. S2. Computational architecture of MOSH. (A) Conceptual schematic illustrating the integration of the volumetric screening modality (eFOV and EDOF super-resolution imaging) and the targeted volumetric modality. The centralized computational platform orchestrates the transition between hardware states, shifting from an active electrically tunable lens (ETL) to an active Z-stage translation. (B) Detailed schematic of the acquisition, reconstruction, and analysis workflow. The computational and hardware workflow is divided into four interdependent zones. Zone 1 illustrates the high-speed 2D image acquisition, utilizing hardware-level TTL triggering and a multi-threaded Producer-Consumer architecture for memory-safe Zarr disk writing. Zone 2 outlines the Dask-accelerated reconstruction process, featuring a human-in-the-loop preview step, macro-level parallel dispatch, pixel reassignment, and flat-field artifact removal. Zone 3 details the AI-driven targeting loop, where Cellpose segmentation and morphological atypia scoring of the cleaned 2D images generate absolute stage coordinates for targeted 3D acquisition. Zone 4 depicts the final visualization stage, where the reconstructed 2D images are stitched into large field-of-view mosaics and computationally converted into virtual H&E images using subtractive optical density transformations.

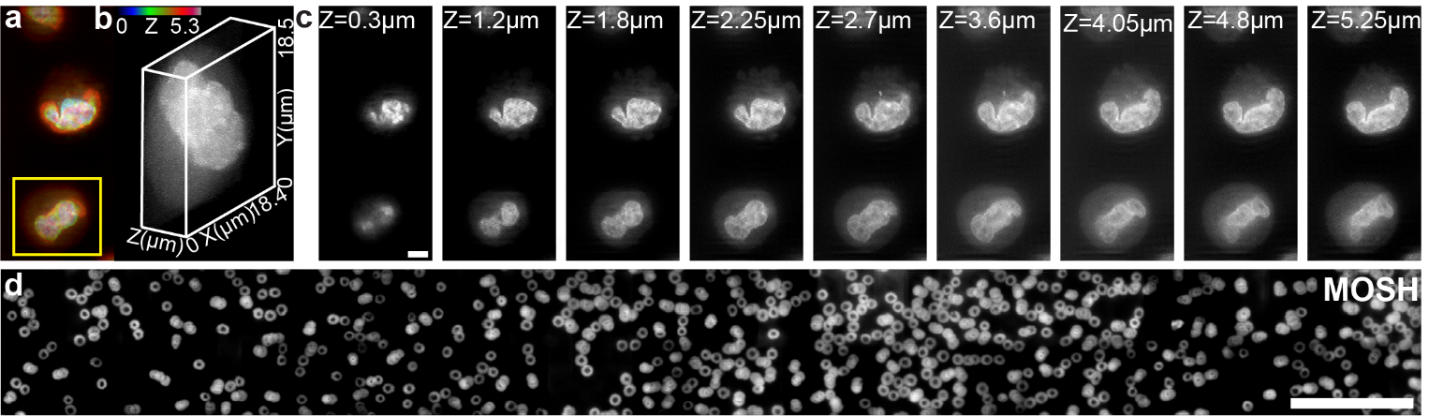

Fig. S3. Cytology and Hematology imaging using MOSH. (A) Depth-color-coded projected image from the volumetric imaging of a K562 nucleus. (B) Zoomed-in volumetric reconstruction of the single nucleus marked in (A). (C) Super-resolution image stack of the nucleus marked in (A), acquired across the depth range. (D) Blood smear imaging across a large FOV and an EDoF of 5 µm. Scale bar: 5 µm (A, C), 100 µm (D).

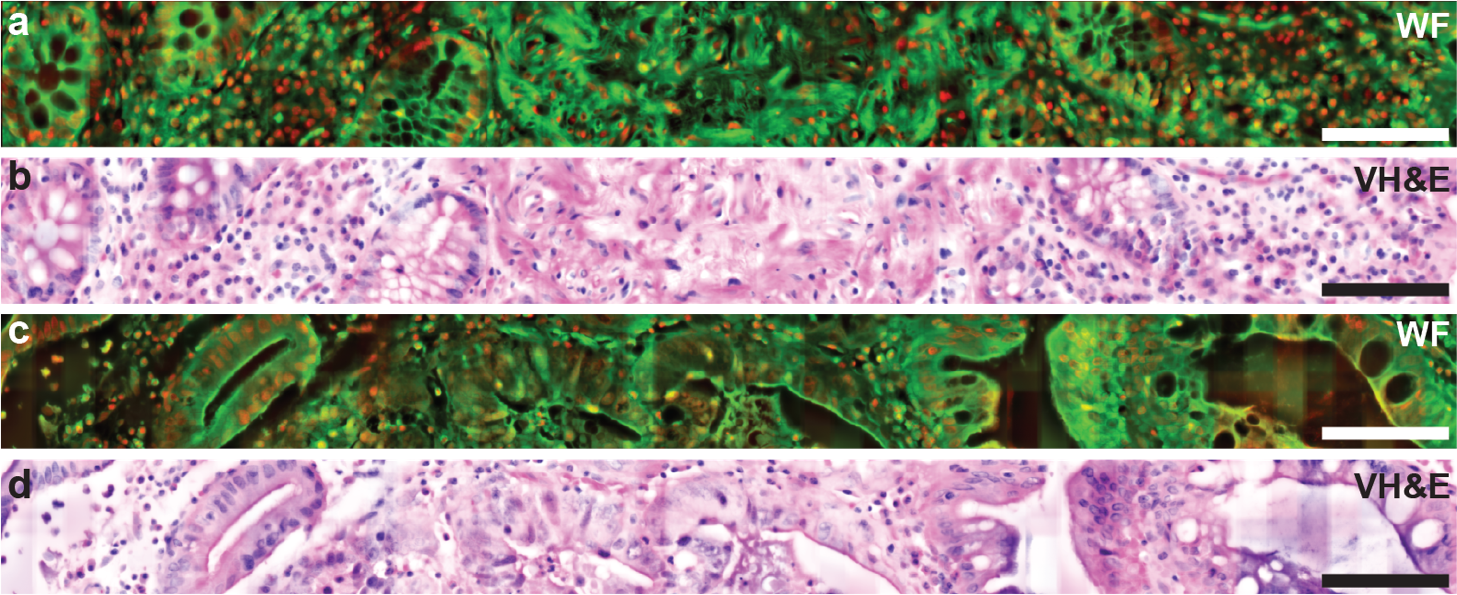

Fig. S4. Wide-field imaging of benign and malignant regions in Stage I colon adenocarcinoma. (A, B) Stacked wide-field (WF) fluorescence image of the benign colon region corresponding to Fig. 3B (A) and its virtual hematoxylin and eosin (vH&E) rendering (B). (C, D) EDoF wide-field (WF) fluorescence image of the malignant colon region corresponding to Fig. 3H (C) and its vH&E rendering (D). The full WF fields are shown to provide the surrounding tissue context for the regions presented in Fig. 3. MOSH acquired a continuous 1.146 mm × 114 µm field of view in (A, C). Scale bars: 100 µm (A-D).

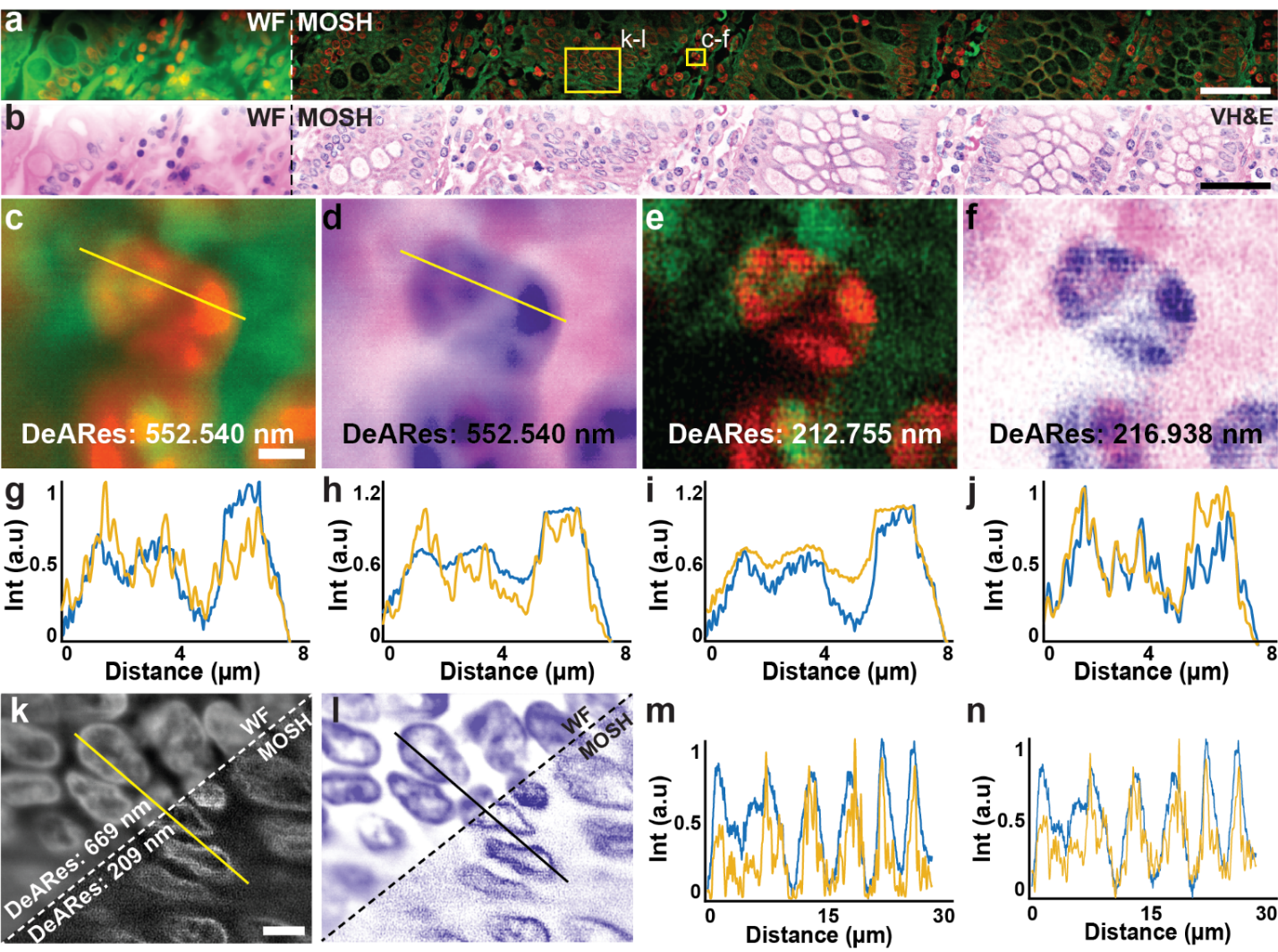

Fig. S5. Improved resolution and optical sectioning with MOSH. (A) Wide-field versus super-resolution fluorescence imaging of stage II colon cancer tissue. The images capture a large field of view (FOV) over a 5 µm extended depth of focus (EDoF). (B) Corresponding wide-field versus super-resolution virtual H&E images derived from the fluorescence images shown in (A). (C) Zoomed-in wide-field image of the region marked in (A). (D) Corresponding zoomed-in wide-field virtual H&E image of the region marked in (A). (E) Zoomed-in MOSH image of the region marked in (A). (F) Corresponding zoomed-in MOSH virtual H&E image of the region marked in (A). (G) Intensity profiles across the lines marked in (C) for the fluorescence images (blue for wide-field, orange for MOSH). (H) Intensity profiles across the lines marked in (D) for the corresponding virtual H&E images (blue for wide-field, orange for MOSH). (I) Intensity profile comparison between the wide-field (orange) and corresponding virtual H&E (blue) images. (J) Intensity profile comparison between the MOSH (orange) and corresponding virtual H&E (blue) images. (K) Zoomed-in nuclear-channel image of the region marked in (A). (L) Corresponding virtual hematoxylin-channel image of (K). (M) Wide-field (blue) versus MOSH (orange) intensity profiles across the line marked in the fluorescence image in (K). (N) Wide-field (blue) versus MOSH (orange) intensity profiles across the line marked in the virtual image in (L). FOV: 909.84 × 64.45 µm (A, B). Scale bars: 50 µm (A, B), 2 µm (C-F), and 5 µm (K, L).

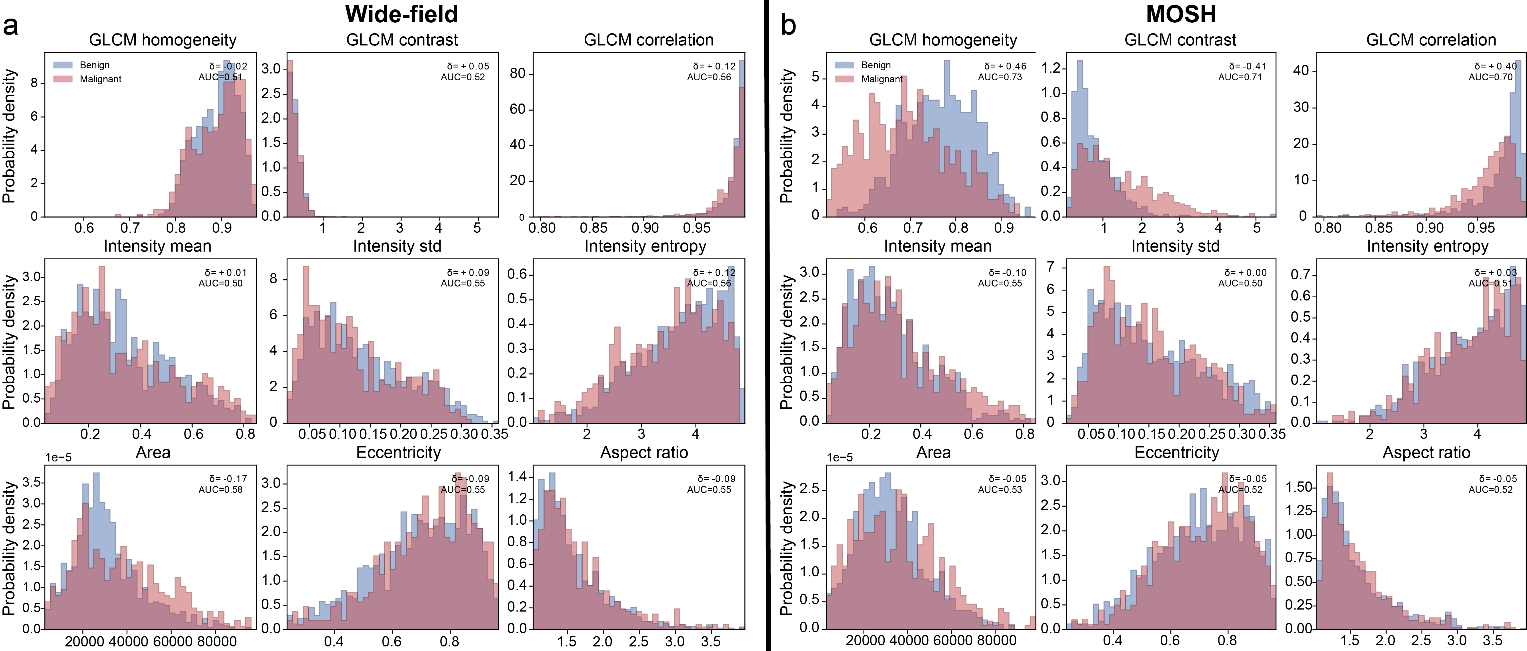

Fig. S6. Widefield Vs. MOSH analysis for benign Vs. Malignant Stage I Colon cancer tissue. (A) Full feature comparison of benign versus malignant nuclei under wide-field (WF) imaging. (B) Full feature comparison of benign versus malignant nuclei under super-resolution (SR) MOSH imaging.

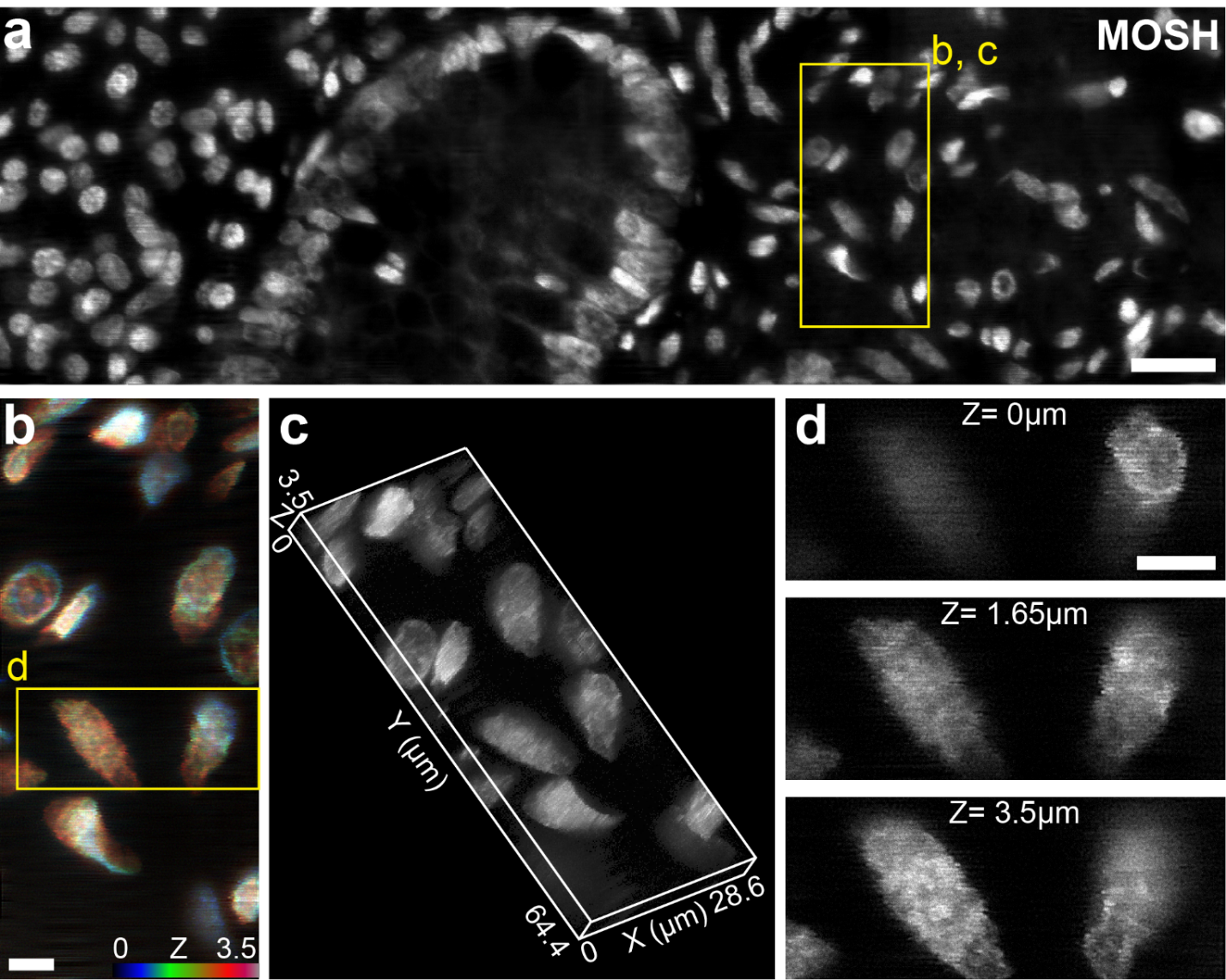

Fig. S7. Multiscale optical super-resolution histology (MOSH) of Stage I human colon cancer tissue. (A) EDoF MOSH nuclear-channel image of a benign region within the tissue sample, demonstrating how the extended depth of focus brings multiple nuclei situated at varying axial depths into simultaneous sharp focus. (B) Depth-color-coded maximum intensity projection (MIP) of the volumetric region highlighted in (A). (C) 3D volume rendering of the region indicated in (A). (D) Sequential super-resolution image stack (optical sections) of the region identified in (B). In contrast to the flattened EDoF projection in (A), these discrete z-planes reveal two distinct nuclei coming into focus at different axial depths. This layer-by-layer structural separation highlights the robust 3D optical sectioning capability of the MOSH platform within thick scattering tissue, isolating internal subnuclear features free from out-of-focus background blur. Scale bars: 20 µm (A) and 5 µm (B, D).

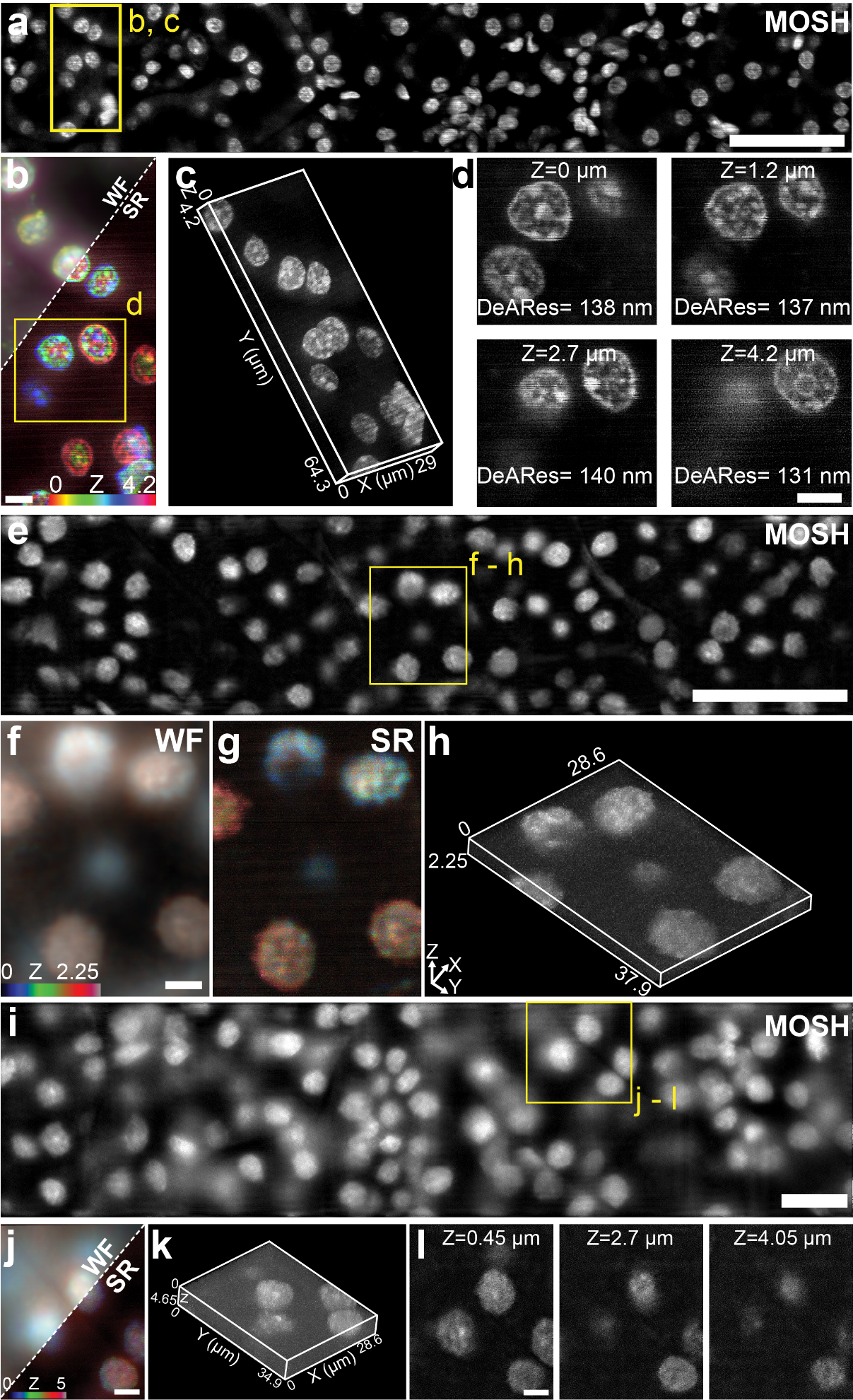

Fig. S8. Additional volumetric and optical-sectioning analysis of mouse kidney tissue across different preparation formats. (A) MOSH nuclear-channel image cropped from the extended field of view in Fig. 4B of an FFPE mouse kidney section. The EDoF image projects fluorescence acquired over the axial range into a single depth-extended representation, maintaining visibility of nuclei located at different axial positions. (B) Depth-color-coded maximum-intensity projection (MIP) of the volumetric region indicated in (A). (C) Three-dimensional rendering of the region indicated in (A). (D) Sequential super-resolution optical sections through the region indicated in (B), showing depth-dependent changes in the visibility and morphology of individual nuclei. (E) MOSH nuclear-channel image cropped from Fig. 4H of an intact perfused mouse kidney punch biopsy acquired over an axial range extending 2.25 µm from the tissue surface. (F, G) Wide-field (F) and super-resolution (G) depth-color-coded MIPs of the volumetric region indicated in (E). (H) Three-dimensional rendering of the region indicated in (E). (I) MOSH nuclear-channel image cropped from Fig. 4I of an intact perfused mouse kidney punch biopsy acquired over an axial range extending 5 µm from the tissue surface. (J) Wide-field and super-resolution depth-color-coded MIPs of the volumetric region indicated in (I). (K) Three-dimensional rendering of the region indicated in (I). (L) Sequential super-resolution optical sections through the nuclei indicated in (I), showing depth-dependent structural changes across the 5 µm acquisition range. Scale bars: 50 µm (A, E), 20 µm (I), and 5 µm (B, D, F, G, J, L).

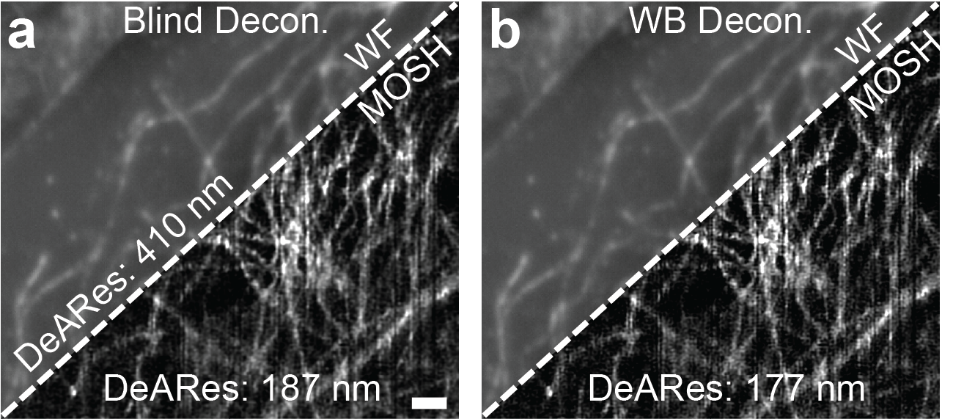

Figure S9: Comparison of deconvolution methods. (A) Widefield Vs. MOSH image with blind deconvolution (5 iterations). (B) Widefield Vs. MOSH image with WB deconvolution (1 iteration). Scale bar: 2 µm (A, B).

Table S1. Comparison of super‑resolution and conventional imaging modalities for histological and pathological tissue analysis. Lateral resolution, area throughput, and maximum tissue thickness are representative values compiled from peer‑reviewed studies; actual performance depends on labeling density, optical configuration, and reconstruction algorithms. “Max tissue thickness” indicates the depth range over which the stated resolution and image quality are typically maintained in practice, not an absolute physical limit. Sample preparation lists the minimal workflow required to achieve the reported performance. Diagnostic output describes the typical data format delivered to pathologists or end users. MOSH denotes the extended depth‑of‑focus, super‑resolved fluorescence and virtual H&E system presented in this work.

| **Modality** | **Lateral Resolution** | **Area Throughput (µm²/s)** | **Demonstrated Tissue Thickness / Imaging Depth*** | **Required Sample Preparation** | **Output** |
| --- | --- | --- | --- | --- | --- |
| Widefield / Confocal (conventional)(*3-5*) | ~200-250 nm (widefield); ~170-200 nm (confocal) | High (widefield); moderate–high (confocal, point‑scan) | ~10–100 µm (confocal, depth limited by scattering) | Standard FFPE or frozen sections | H&E, IF, brightfield / fluorescence |
| Clinical SIM(*6, 7*) | ~1.3 µm | ~3.3×106 | < 5 µm,  Thick intact tissue demonstrated | Physical microtomy (FFPE) | Pseudo‑color / H&E overlay |
| Expansion Microscopy (ExM)(*8*) | ~60-70 nm (effective) | Very Low (≪10²) | Up to hundreds of µm pre expansion, (protocol‑dependent) | Proteolytic digestion, gel embedding, expansion | Fluorescence |
| SMLM / STORM on Tissue(*3, 9*) | ~20 nm | < 1 (extremely slow) | < 2–3 µm (practical depth) | Physical microtomy, blinking buffers | Fluorescence |
| Commercial SIM (Tissue)(*10*) | ~100-120 nm | ~10²–10³ (configuration‑dependent) | ~0.3–1 µm (optimal), up to ~20 µm possible with degraded resolution | Physical microtomy (FFPE/frozen) | Fluorescence |
| **MOSH (This Work)** | **~120-150 nm** | **~715 (with 20 µm EDoF)** | **~20 µm EDoF (unsectioned)** | **Standard FFPE or fresh intact tissue** | **Fluorescence & SR vH&E** |

Table S2: List of components used in the MOSH system.

| **PARTS** | **SOURCE** | **IDENTIFIER** |
| --- | --- | --- |
| Microscope frame | Nikon | Eclipse Ti2-U |
| Laser 1 | Edmund Optics | Coherent, OBIS 488 nm LX 100 mW |
| Laser 2 | Edmund Optics | Coherent, OBIS 647 nm LX 120 mW |
| High-precision motorized stage | Applied Scientific Instrumentation | MS-2000FT |
| High-precision piezo Z stage | Mad City Labs | Nano Drive |
| Electrotuneable lens | Optotune | EL-10-30-C-VIS-LD-MV |
| Electrotuneable lens driver | Optotune | Lens driver 4 |
| Dichroic mirror | Chroma | ZT488/640rpc-UF1 |
| Emission filter | Chroma | ZET488/640m |
| Telescope lens 1 | Thorlabs | AC254-030-A-ML |
| Telescope lens 2 | Thorlabs | AC254-500-A-ML |
| Collimator lens | Thorlabs | AC254-100-A-ML |
| Relay lens ×2 | Thorlabs | AC254-030-A-ML |
| sCMOS camera | Tucsen | Dhyana 400BSI V3 |
| Microlens array | RPC Photonics | MLA-S100-f4-A-R1 |

Table S3: List of reagents and consumables

| **ITEM** | **SOURCE** | **IDENTIFIER** |
| --- | --- | --- |
| Poly-L-lysine (0.01%) | Thermo Fisher Scientific | NC3834048 |
| Paraformaldehyde (PFA, 4%) | Electron Microscopy Sciences | 15710 |
| PBS | Corning | 21-040-CV |
| DRAQ5 | Thermo Fisher Scientific | 62251 |
| Eosin Y | Thermo Fisher Scientific | 152880250 |
| Methanol (Absolute) | Thermo Fisher Scientific | 031721.AP |
| Ethanol | Thermo Fisher Scientific | P111WORLD200PL05 |
| Glycerol (100%) | Thermo Fisher Scientific | 332031000 |
| Xylene | Thermo Fisher Scientific | X5-4 |
| Imaging Dish 35 mm | World Precision Instruments | FD35-100 |
| Dermal punch biopsy (Model MB) | MedBlades |  |
| Glass microscope slides | VWR international | 16004-370 |
| Coverslips (No. 1) | VWR international | 16004-308 |
